# Restriction enzyme selection dictates detection range sensitivity in chromatin conformation capture-based variant-to-gene mapping approaches

**DOI:** 10.1101/2020.12.15.422932

**Authors:** Chun Su, Matthew C. Pahl, Struan F. A. Grant, Andrew D. Wells

## Abstract

Promoter-focused chromatin conformation techniques directly detect interactions between gene promoters and distal genomic sequences, providing structural information relevant to gene regulation without the excessive non-genic architectural data generated by full scale Hi-C. 3D promoter ‘interactome’ maps are crucial for understanding how epigenomic features like histone modifications and open chromatin, or genetic variants identified in genome wide association studies (GWAS), contribute to biological function. However, variation in sensitivity between such promoter-focused methods, principally due to restriction enzyme selection, has not been systematically assessed. Here, we performed a head-to-head comparison between promoter capture C (PCC) and promoter capture Hi-C (PCHiC) with the respective 4 cutters DpnII and MboI versus 6 cutter HindIII datasets from the same five cell types. While PCHiC generally produces a higher signal to noise ratio for significant interactions in comparison to PCC, we show that DpnII/MboI detects more proximal interactions and shows little overlap with the HindIII detection range. Promoter-interacting genomic regions mapped by 4-cutters are more enriched for regulatory features and disease-associated genetic variation than 6-cutters maps, suggesting that high-resolution maps better capture gene regulatory architectures than do lower resolution approaches.

---

Chromatin conformation capture (3C) based methods combined with high-throughput deep sequencing are increasingly used to detect direct interactions between gene promoters and distal genomic sequences. These techniques specifically enrich for promoter-containing DNA fragments using complementary RNA oligomers, increasing the resolution and cost-efficiency of detecting gene interactions compared to conventional Hi-C, which generates interaction data across the whole, largely intergenic, genome. The resulting promoter contact maps are widely used to assign distal regulatory elements to their target genes for a better understanding of global regulation of gene expression^1,2^, and to connect human disease-associated variants to their effector gene promoters following genome-wide association studies (GWAS)^3–5^.

Promoter capture C (PCC) is applied directly to 3C libraries, while promoter capture Hi-C (PCHiC) involves an additional streptavidin-biotin pulldown step before promoter capture hybridization to enrich for genuine ligation products and increase the efficiency of sequencing valid hybrid reads^6^. To compare the results of different promoter capture techniques, we selected five distinct cell types with datasets available for both PCC and PCHiC^1^ (**Supplemental Table 1**). As previously estimated^7^, PCHiC read usage was 2~3 times more efficient than PCC, as measured by the ratio of valid read pairs to mapped read pairs (**Supplemental Table 1**). More dramatically, PCC data was enriched for valid read pairs that mapped to a shorter range (**Supplemental Figure 1 A**), which may reflect the contribution of undigested fragments instead of *bona fide* ligation products. The retention of partially digested fragments in single-fragment analyses skews toward a higher percentage of short-range significant interactions in PCC samples (**Supplemental Figure 1 B**), which may be artefactual. To mitigate the influence of partial digestion in PCC, we concatenated neighboring fragments *in silico* by binning valid reads from 4 fragments into one larger fragment, effectively filtering out interactions potentially called due to partial digestion. This approach not only attenuated over-representation of short-range interactions in the PCC data (**Supplemental Figure 1B**), but also increased the supporting read count per fragment to detect longer-distance interactions.

Another key variation in library preparation is the choice of restriction enzyme. The widely used restriction enzymes include HindIII with a 6 bp recognition site^6,1,8,9^ and DpnII or MboI with a 4 bp recognition site^4,10,3,11^. A computational DpnII or MboI restriction digest of the human genome predicts fragments 7 to 9 times smaller than a HindIII digest (**Supplemental Figure 2 A**), leading to the potential for higher-resolution interaction maps. In addition, the smaller fragment size of DpnII and MboI leads to more precise annotation, pinpointing more than 75% of captured fragments (baits) to just one transcriptional start site (TSS), and resulting in a median fragment size of 265 bp in one fragment resolution (**Supplemental Table 2**). This resolution equates to approximately two nucleosomes, or the size of a single chromatin accessible region^12^, an important consideration if one wants to integrate with ATAC-seq or ChIP-seq datasets. An example of this is shown at the *GRAP2* and *ENTHD1* genes in naïve CD4 T cells, where DpnII PCC can resolve the interactomes of each promoter, but HindIII PCHiC cannot (**Supplemental Figure 2 B**).

CHiCAGO is the most commonly used software for calling interactions for both PCC and PCHiC^13^. With minor differences in parameter settings which take the fragment size difference into account (see Methods), PCC and PCHiC, regardless of resolution, generate a comparable number of total interactions, *cis-trans* interaction ratios, and bait-to-bait interaction ratios (**Supplemental Table 3**). However, with the higher captured read-pair input, each significant interaction is supported with a higher average number of reads in HindIII PCHiC (**Supplemental Figure 3**), leading to a higher signal-to-noise ratio.

One caveat in using CHiCAGO calls is that bait-to-bait interactions are not always called as “significant” at both baits, even though physically they must be bi-directional. This results from CHiCAGO implementing bait-specific and counterpart-specific (other-end) factor scaling for Brownian background estimation. The internal consistency between bait-to-bait interactions can provide an intra-library metric for reproducibility. Compared to HindIII PCHiC, we observed a significantly higher bi-directional (reproducible) bait-to-bait interaction ratio for DpnII PCC (**Supplemental Figure 4 A, Supplemental Table 3**). This likely results from less dispersed bait-specific and other-end-specific scaling factors in the DpnII PCC design (**Supplemental Figure 4 B**), and suggests that DpnII bait design results in more uniform hybridization across baits.

While different capture techniques and restriction enzymes yield roughly comparable quality control metrics as outlined above, these approaches yield drastically different results with regard to the detection range of promoter interactions. The detection ranges of MboI PCHiC, DpnII PCC and HindIII PCHiC are all consistent with the size of mammalian topologically associating domains (TADs), which range from 100kb to 5Mb with an average of ~1Mb^14^. However, regardless of different capture techniques, while the HindIII detection range was between 150kb and ~ 2Mb, DpnII-based and MboI-based 4-cutter libraries detect many more proximal interactions (**Figure 1 A** and **B**). Ninety percent of interactions detected by DpnII PCC were concentrated between 500 bp and 500 kb (single fragment call: 500bp ~ 300kb; 4 fragment call: 2kb ~ 500kb), and the MboI PCHiC detection distance was distributed from 6kp to 500kb (**Figure 1 A** and **B**). This difference in range resulted in only 10 to 40% of detected interactions overlapping between HindIII and the 4-cutters, depending upon fragment resolution, library quality and Chicago settings (**Supplemental Table 4**). We also compared the two methods at the gene/promoter level, by comparing only baits harboring the same gene TSS to account for differences in bait design and promoter annotation between the two approaches (**Supplemental Table 5**). Approximately 65% of genes were engaged in at least one significant interaction with a distal region in both the DpnII PCC and HindIII PCHiC libraries, with twice as many DpnII-specific interacting genes called compared to HindIII (**Figure 1 C**, left panel). The set of interactive genes detected by MboI PCHiC vs. HindIII PCHiC is even less similar, with only 43% of genes having at least one significant distal interaction in both the MboI PCHiC and HindIII PCHiC CM libraries (**Figure 1 C**, right panel). To further explore the drivers behind the resolution-specific genes, we compared the interaction distance for 4-cutter-specific interacting genes to those common to both cutters, and found that 4-cutter-specific interacting genes involved contacts at significantly shorter distances (**Figure 1 D**, two-tailed Wilcoxon sum rank test). This suggests that the 4-cutter methods detect contacts that are too promoter-proximal to be detected in HindIII PCHiC datasets. Indeed, we could find a myriad of sub-150kb promoter contacts revealed by DpnII-resolution PCC that were not detected by HindIII PCHiC. For example, DpnII PCC detected dozens of interactions within 150 kb of the baited *SERBP1* promoter in naïve B cells, including open regions residing in introns of *IL12RB2.* However, those interactions were not detected in HindIII PCHiC (**Figure 1E**). Conversely, DpnII PCC with 4-fragment binning failed to detect very long-range interactions detected with HindIII (>500kbp). However, by increasing fragment concatenation levels further (6-, 8-fragments, etc), the DpnII method has the potential to complement long-range interaction detection.

**Figure 1.**
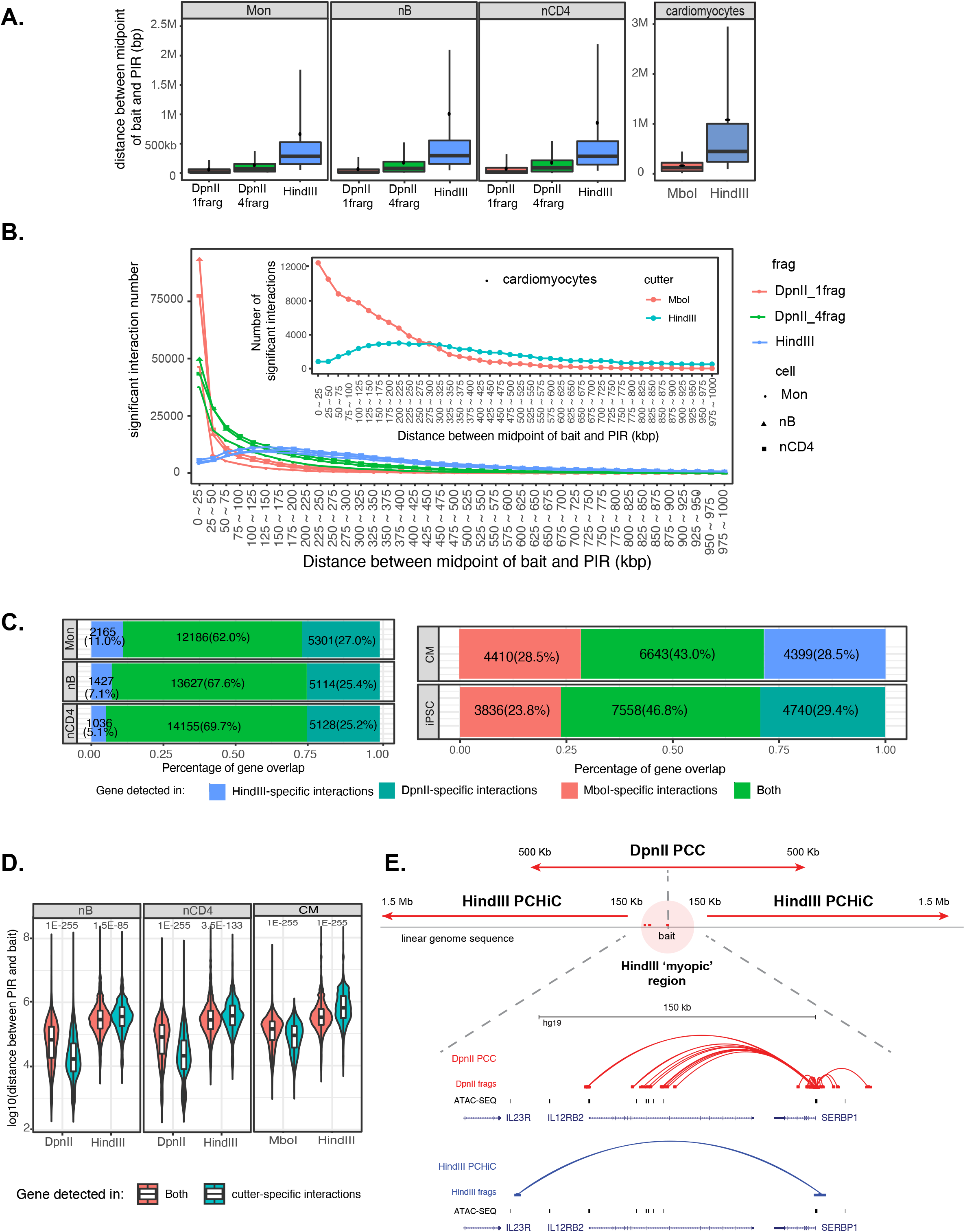
Interaction range detection difference results in low overlap between DpnII and HindIII. **A.** Distance ranges of intra-chromosomal interaction In DpnII PCC vs. HindIII PCHiC (panels 1-3) and MboI PCHiC vs. HindIII PCHiC (panel 4). The distance between mid-points of bait fragment and PIR was plotted in boxplot for significant cis-interactions (CHiCAGO score > 5). The upper whisker, upper hinge, middle line, middle dot, down hinge and down whisker indicate 95%, 75%, median, mean, 25% and 5% percentile. **B**. The number of significant cis-interaction across different distance between bait and PIR. The significant cis-interactions in DpnII PCC vs. HindIII PCHiC were grouped based the binned distance (250k intervals) between mid-points of bait fragment and PIR. lnteraction number within each group was plotted. The inset depicts comparative distance interaction distributions of MboI PCHiC vs. HindIII PCHiC. **C.** Venn diagram of genes annotated in DpnII PCC, MboI PCHiC, and HindIII PCHiC datasets. Genes were annotated when the bait of a significant interaction harbors its TSS. Regardless of PIR overlaps, shared (light green), DpnII-specific (dark green), MboI-specific (red), and HindIII-specific (blue) genes were determined based on whether corresponding baits have at least one significant distal interaction in both DpnII and HindIII, MboI and HindIII, or DpnII and MboI datasets. **D.** Distance difference between interactions involving cutter-specific and shared genes. The interaction distance distribution measured by each method (x-axis) was calculated for cutter-specific gene sets (blue) or shared gene sets (red). The labeled p-value was determined by two-tailed wilcoxon rank sum test. **E**. Example of *SERBP1* interacting region range difference between DpnII PCC and HindIII PCHiC in naïve B cells. The DpnII and HindIII baits for *SERBP1* were overlapped. Only significant interactions (CHiCAGO score > 5) were shown with arcs in either DpnII PCC (red) and HindIII PCHiC (blue).

To further investigate the regulatory significance of the difference in detection range between 4-cutters and HindIII, we overlapped the promoter-interacting genomic regions with regulatory features determined by ATAC-seq for open chromatin and ChIP-seq for histone marks (**Supplemental Table 6**) in the respective cell types. Promoter-interacting genomic regions detected by 4-cutter PCC and PCHiC were significantly more enriched for active promoter marks (H3K4me3), enhancer marks (H3K27ac and H3K4me1), and regions of open chromatin (ATAC-seq, **Figure 2 A** and **B**) compared to HindIII PCHiC, but less enriched for the H3K36me3 mark that is enriched at the 3’ ends of transcriptionally elongating genes and contributes to the composition of heterochromatin^15^. We observed the same enrichment of active chromatin signatures by DpnII in both blood-derived and tonsil-derived naïve B cells (**Figure 2 A)**, suggesting that this enrichment is not an artifact of donor or tissue of origin, but reflects a real difference between resolutions. Furthermore, to remove the influence of artefactual interactions due to partial digestion in PCC, we filtered out interactions called at less than 4 fragments away from a bait. Importantly, removal of these supra-proximal interactions from the 1 fragment resolution PCC datasets did not impact the enhanced enrichment of regulatory regions in DpnII PCC libraries compared to Hindll PCHiC libraries, indicating that partial digestion artifacts in PCC libraries do not significantly impair the ability of this technique to capture interactions between gene promoters and regulatory chromatin elements (**Supplemental figure 5**). Overall, these results suggest that 4-cutter libraries detect functionally relevant regulatory regions than the lower resolution HindIII-based maps.

**Figure 2.**
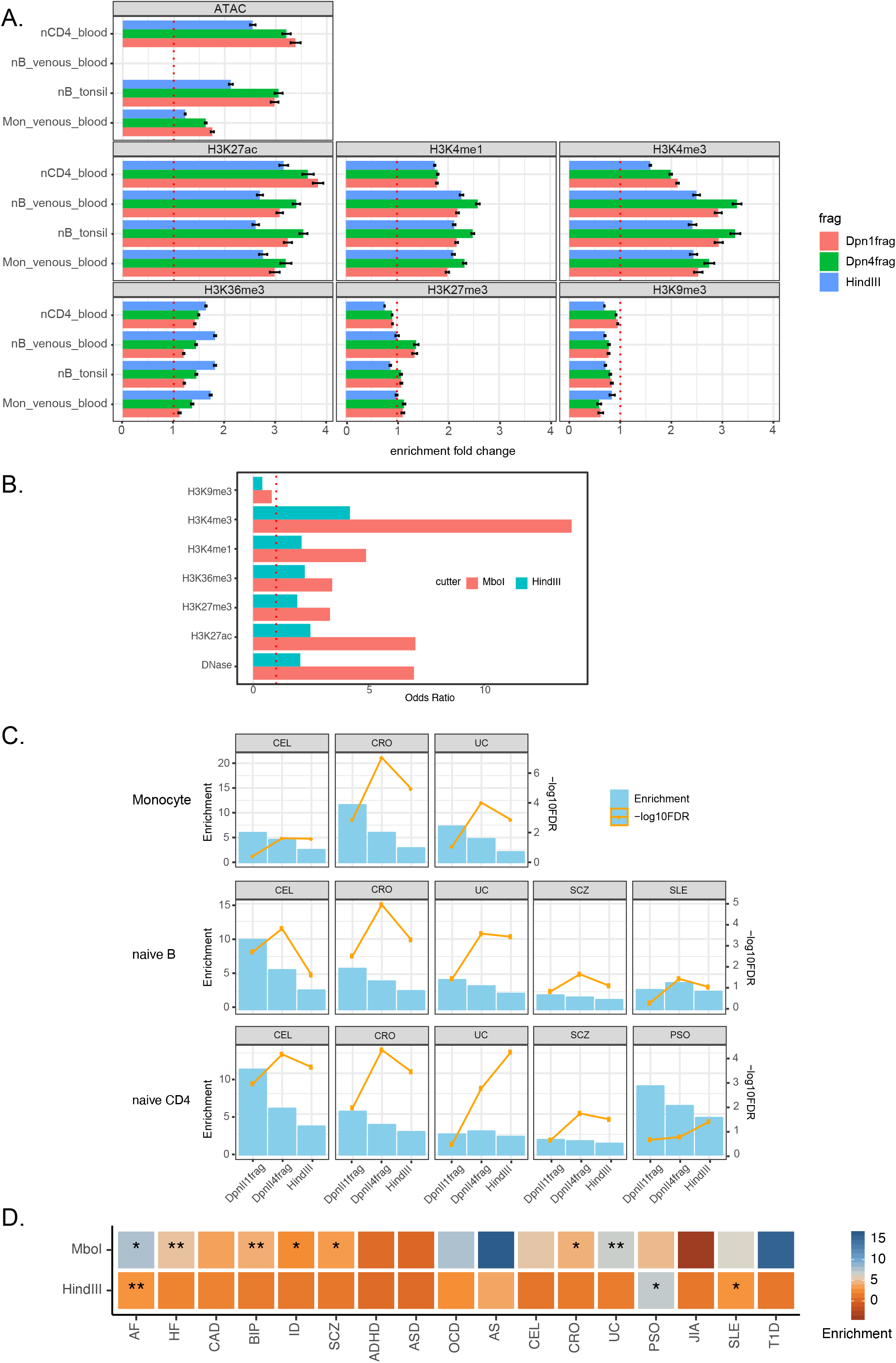
Enrichment of regulatory DNA and heritability of immune and psychiatric traits by PIRs from immune cells. **A**. Enrichment of regulatory regions by promoter-interacting regions. The number of PIR overlapped with open chromatin and histone binding regions were calculated in both significant interactions and randomly distance-matched non-significant interactions. The enrichment fold change is measure by the ratio of PIR number between significant and non-significant interactions. Mean□±□95% CI is depicted across 100 draws of non-significant interactions. **B**. Enrichment of regulatory regions by promoter-interacting regions in cardiomyocytes. The odds ratio is determined by Locus Overlap Analysis (LOLA)^20^ for both cutters independently. It compares PIRs identified by significant interactions to PIRs identified by non-significant interactions using Fisher’s exact test. **C**. Heritability enrichment of autoimmune and psychiatric disorders by PIRs in immune cells. Partitioned LD score regression was performed independently in three immune cell types (Monocyte, naïve B and CD4 T cells) across 15 GWAS studies and PIRs detected by different cutters. Enrichment of heritability (blue) was measured in PIRs compared to baseline, and p-value was adjusted with FDR (orange). **D**. Heritability enrichment of GWAS traits by PIRs in cardiomyocytes. Partitioned LD score regression was performed independently across 18 GWAS studies and PIRs detected by different cutters. Enrichment of heritability was measured in PIRs compared to baseline and labeled with heatmap. The significant enrichment was indicated by stars (* *P* < 0.05, ** *P* < 0.01). Details can be found in **Supplemental Table 7**.

One important application of promoter capture data is to discover the effector genes that may be regulated by disease-associated causal variants in relevant tissues, *i.e.* variant-to-gene mapping. To compare the efficiency of causal variant detection by 4-cutter- and 6-cutter-based approaches, we performed linkage disequilibrium score regression (LDSC) to estimate the heritability content of PIRs in three immune cell types across eight autoimmune and seven psychiatric arbitrarily selected GWAS datasets, and PIRs in cardiomyocytes across three cardiac-related traits (**Supplemental Table 6**). In general, variants associated with Crohn’s disease, ulcerative colitis, and celiac disease were enriched in all three immune cell types, schizophrenia variants were modestly enriched in T and B lymphocytes (**Figure 2 C**, FDR<0.05, **Supplemental Table 7**), and variants associated with atrial fibrillation and heart failure were highly enriched in cardiomyocytes (**Figure 2 D**, FDR<0.05, **Supplemental Table 7**). Compared to promoter interacting genomic regions derived from HindIII PCHiC, the shorter-range regions detected by DpnII PCC and MboI PCHiC were more enriched for heritability of the corresponding traits (**Figure 2 C and D**). This same trend between distance and heritability could be observed within the HindIII PCHiC datasets themselves, as disease heritability was enriched in the set of HindIII-based interacting regions located less than 500kb away from promoters, but no significant enrichment was detected for regions beyond 500kb (**Supplemental Figure 6 A**). Finally, as also observed with feature enrichment, the enhanced heritability enrichment of cell type-relevant traits by high-resolution PIRs was not impacted if potentially artefactual interactions due to partial digestion is filtered from the PCC datasets (**Supplemental Figure 6 B**). Together, these results demonstrate that high resolution DpnII and MboI maps capture disease-relevant promoter interactions to a greater extent than HindIII-based maps, and importantly, imply that disease-causal variants are most commonly located less than 500kb from the promoter of their effector gene.

Overall, we conclude that distinct promoter-chromatin conformation capture techniques and restriction enzymes detect distinct promoter interactomes in the same cell types, owing largely to very different interaction range distributions. HindIII PCHiC maps excel at detecting interactions >150kb to 2Mb from promoters, but potentially bias investigators away from genes nearest GWAS signals. Conversely, higher resolution PCC and PCHiC maps are more sensitive at detecting highly interactive gene regulatory architectures built from active chromatin regions located relatively proximal (<150kb) to gene promoters. High-resolution gene regulatory architectures are also enriched for variants that are strongly associated with relevant common complex traits in given cell types. Moreover, the power of 4-cutter-based capture techniques to detect longer-range interactions can be increased through *in silico* binning of reads from neighboring fragments, which also eliminates partial digestion artifacts inherent to PCC libraries. Therefore, publicly available promoter capture datasets generated with different restriction enzymes should not be considered as comparable, even from the same cell types, and investigators should carefully consider which promoter capture approaches or publicly available datasets best fit the goals of their research.

## Methods

### Promoter capture bait design

Bait design for HindIII PCHiC datasets was downloaded from Open Science Framework (https://osf.io/u8tzp), while MboI PCHiC bait design was derived from **Supplemental Table 9** by Montefiori et al^10^. DpnII PCC designs contains both 1-fragment and 4-fragment resolution and were obtained from Array Express (<u>E-MTAB-6862</u>)^3^. To make bait design comparable across the studies, all baits were re-annotated to genes using bait coordinates overlapping with transcription start site (TSS) from Ensembl v73. The bait design comparison is summarized in **Supplemental Table 2**.

### Promoter capture C pre-processing and interaction calling

DpnII PCC libraries from 3 to 4 donors for monocytes, naïve CD4+, naïve B and iPSC cells were pre-processed using the HiCUP pipeline (v0.5.9)^16^, with bowtie2 as aligner and hg19 as the reference genome. Unique captured read pairs from all baited promoters were used for significant promoter interaction calling. Significant promoter interactions at 1-fragment resolution were called using CHiCAGO^13^ (v1.1.8) with default parameters except for binsize set to 2500. Significant interactions at 4-DpnII fragment resolution were also called using CHiCAGO with artificial baitmap and rmap files in which DpnII fragments were concatenated *in silico* into 4 consecutive fragments using default parameters except for removeAdjacent set to False and binsize set to 10000. Interactions with a CHiCAGO score > 5 in either 1-fragment or 4-fragment resolution were considered as significant interactions. The significant interactions were finally converted to *.ibed* format in which each line represents a physical interaction between fragments.

HindIII PCHiC immune datasets were also pre-processed using HICUP by Javierre et al^1^. The summary statistics were extracted from the original paper (**Supplemental Table 1**). CHiCAGO R objects were downloaded from Open Science Framework (https://osf.io/u8tzp) and significant interactions were exported using function “exportResults” with CHiCAGO score > 5. The significant interactions for HindIII PCHiC cardiomyocytes were identified by Choy et al^17^ for each replicate independently. The reproducible interactions that are significant in at least two replicates were used for comparison.

MboI PCHiC datasets (cardiomyocytes and iPSC) were pre-processed using HiCUP by Montefiori et al^10^. The summary statistics were extracted from the original paper (**Supplemental Table 1**). The significant interactions for HindIII PCHiC were identified for each replicate independently by original paper. The reproducible interactions that are significant in at least two replicates were used for further comparison.

The significant interactions from HindIII PCHiC, DpnII PCC and MboI PCHiC can be viewed at <u>UCSC browser</u>.

### ATAC-seq and histone mark peaks

ATAC-seq peaks were called using the ENCODE ATAC-seq pipeline (https://www.encodeproject.org/atac-seq/). Briefly, pair-end reads from all replicates for each cell type were aligned to hg19 genome using bowtie2, and duplicate reads were removed from the alignment. Aligned tags were generated by modifying the reads alignment by offsetting +4bp for all the reads aligned to the forward strand, and −5bp for all the reads aligned to the reverse strand. Narrow peaks were called independently for pooled replicates for each cell type using macs2 (-p 0.01 --nomodel --shift −75 --extsize 150 -B --SPMR --keep-dup all --call-summits) and ENCODE blacklist regions (wgEncodeDacMapabilityConsensusExcludable.bed.gz) were removed from called peaks. The reference peaks for each cell type were merged peaks present in at least half of replicates.

Histone mark peaks were downloaded as processed bed files from The BLUEPRINT Data Analysis Portal^18^ **(Supplemental Table 6).** The reference peaks for each cell type and tissue were obtained by merging replicates and selecting peaks present in at least half of replicates. The hg38 coordinates were finally converted to hg19 coordinates using UCSC liftOver.

### Feature enrichment

When Chicago objects were available (Naïve CD4+, Naïve B and Monocytes), the feature enrichment was performed independently for each cell type and resolution, using “peakEnrichment4Features” function from CHiCAGO (v1.1.8) and plotted with ggplot2. For the samples without Chicago objects (cardiomyocetes), PIRs identified by significant interactions was compared to PIRs identified by non-significant interactions by overlapping with chromatin features in corresponding cell type using Locus Overlap Analysis (LOLA)^20^. The odds ratio is determined by Fisher’s exact test for both cutters independently.

### Partitioned heritability LD score regression enrichment analysis

Partitioned heritability LD Score Regression^19^ (v1.0.0) was used to identify heritability enrichment with GWAS summary statistics and PIRs characterized by Chicago. The baseline analysis was performed using LDSCORE data (https://data.broadinstitute.org/alkesgroup/LDSCORE) with LD scores, regression weights, and allele frequencies from 1000G Phase1. The summary statistics were downloaded using the links and reference provided in **Supplemental Table 6**. The Partitioned LD score regression annotations were generated using the coordinates of non-bait PIRs independently for each cell type and cutters. Finally, the cell-type-specific partitioned LD scores were compared to baseline LD scores to measure enrichment fold change and enrichment p-value were adjusted using FDR across all comparisons. The details of the LDSC enrichment results are provided in Supplemental Table 7.

## Supporting information

supfigure1

supfigure2

supfigure3

supfigure4

supfigure5

supfigure6

---

**Supplemental Figure 1**. Difference between PCC and PCHiC. The percentage of cis-capture read pair count (**A**) and cis significant interaction number (**B**) across different fragment intervals between bait and PIR. PCC (red) and PCHiC (green) libraries were both prepared by 4-cutters for iPSC cells. *In silico* 4-fragment resolution is indicated by triangle.

**Supplemental Figure 2** Resolution comparison between DpnII/MboI and HindIII design. **A.** Fragment length of DpnII/MboI and HindIII by *in silico* digesting hg19 genome. DpnII and MboI recognized the same sequences (^GATC) while HindIII recognize 6-bp sequence (A^AGCTT). Fragment length was plotted in boxplot for all fragments (left) and bait fragments (right). The upper whisker, upper hinge, middle line, middle dot, down hinge and down whisker indicate 95% percentile, 75% percentile, median, mean, 25% percentile and 5% percentile. **B**. Example of bait annotation precision difference between DpnII PCC and HindIII PCHiC at *ENTHD1* and *GRAP2.* Black bars indicate the baits. Two baits in DpnII (red arcs) design represents *ENTHD1* and *GRAP2* independently while these two genes are baited by the same fragment in HindIII (blue arcs) design.

**Supplemental Figure 3**. The normalized read count of significant cis-interaction across distance between bait and PIR. The significant cis-interactions were grouped based the binned distance (250k intervals) between mid-points of bait fragment and PIR. The reads were normalized by total “reference” read count for each cell. The median normalized read count within each group were plotted.

**Supplemental Figure 4**. Bi-directional (reproducible) bait-to-bait interactions detected by DpnII PCC and HindIII promoter capture HiC. **A**. Bait-to-bait interaction self-reproducibility. The self-reproducibility was measured by ratio between bi-directional and all bait-to-bait significant interactions (CHICAGO ≥ 5). **B**. Distribution of bait-specific and PIR-specific scaling factors. The scaling factors were extracted from CHICAGO R objects across all interactions. The average of scaling factors per fragment among all cell types are plotted for distribution.

**Supplemental Figure 5.** Regulatory feature enrichment of PIRs by filtered out interactions called at less than 4 fragments away from a bait. The number of PIR overlapped with open chromatin and histone binding regions were calculated in both significant interactions and randomly distance-matched non-significant interactions. The enrichment fold change is measure by the ratio of PIR number between significant and non-significant interactions. Mean□±□95% CI is depicted across 100 draws of non-significant interactions.

**Supplemental Figure 6**. LDSC enrichment in CEL, CRO and UC in immune cells. **A**. Heritability of autoimmune traits is enriched at PIRs within 500kbp from gene promoter. Partitioned LD score regression was performed independently for PIRs within and beyond 500kbp from HindIII PCHiC data sets. Enrichment of heritability (blue bar) was measured in PIRs of given distance compared to baseline, and p-value was adjusted with FDR (orange point). **B**. Higher heritability enrichment of PIRs in DpnII PCC is not impacted by artefactual interactions from the partial digestions. Partitioned LD score regression was performed independently for a set of DpnII PCC PIRs from which interaction with less than 2 to 20 fragments away were removed. The heritability enrichment of HindIII PCHiC PIRs is indicated by black dashed lines.

## SUPPLEMENTAL TABLES

**Supplemental Table 1: Hicup pre-processing of HindIII PCHiC, Dpn II PCC and MboI PCHiC libraries used in this study.**

**Supplemental Table 2: Bait design and annotation of HindIII PCHiC, Dpn II PCC and MboI PCHiC**

**Supplemental Table 3: Comparison of total and significant interaction calls from CHiCAGO.**

**Supplemental Table 4: The pairwise comparison of significant cis interactions among HindIII PCHiC, Dpn II PCC and MboI PCHiC libraries within the same cell type.**

**Supplemental Table 5: The pairwise comparison of identified gene-PIR interactions among HindIII PCHiC, Dpn II PCC and MboI PCHiC libraries within the same cell type.**

**Supplemental Table 6: External resource for chromatin feature and GWAS summary statistic.**

**Supplemental Table 7: Partitioned LD score regression of PIRs on GWAS traits**

