## Supplementary figures and images for "Restriction enzyme selection dictates detection range sensitivity in chromatin conformation capture-based variant-to-gene mapping approaches"

### supfigure1

##### Supplemental Figure 1

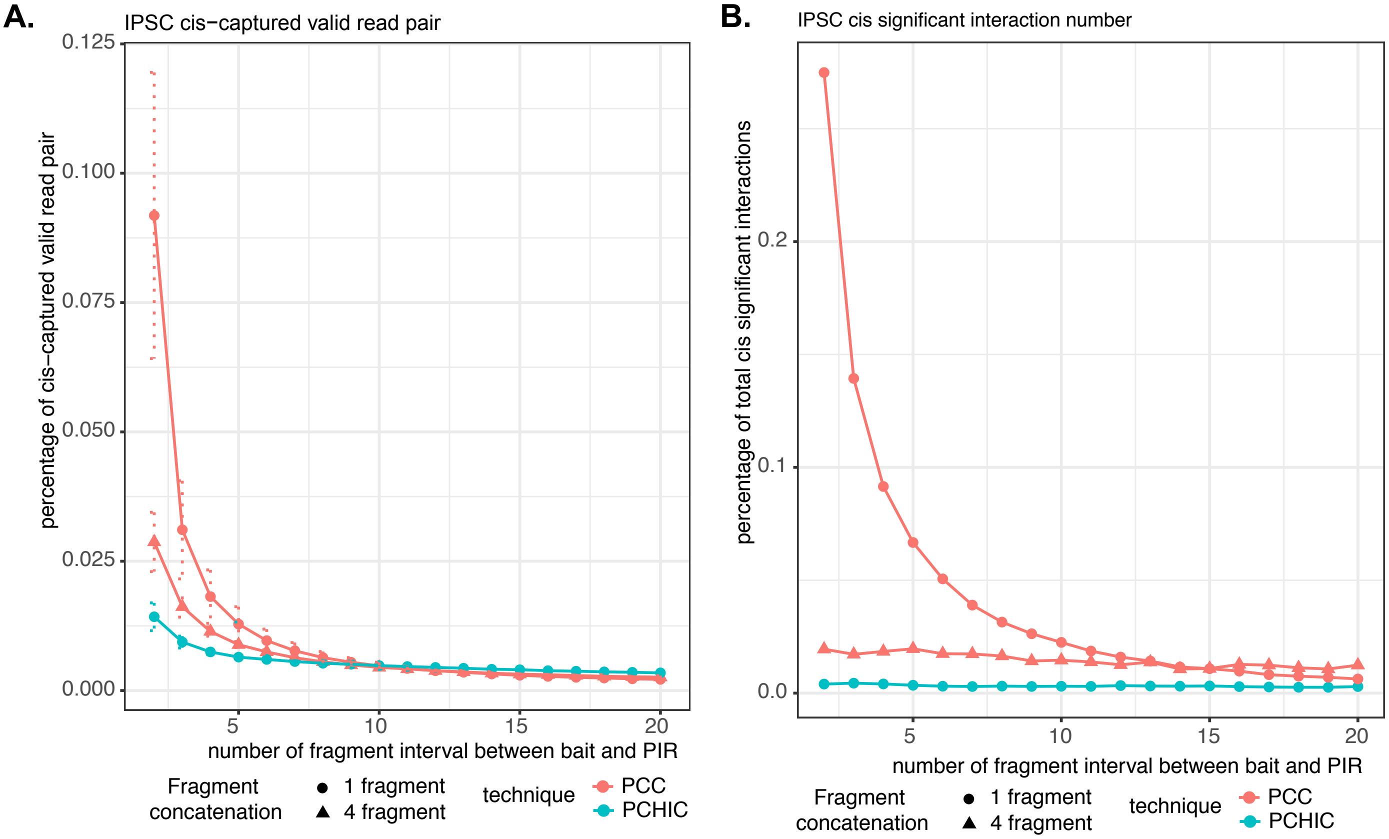

### supfigure2

##### Supplemental Figure 2

**A.**

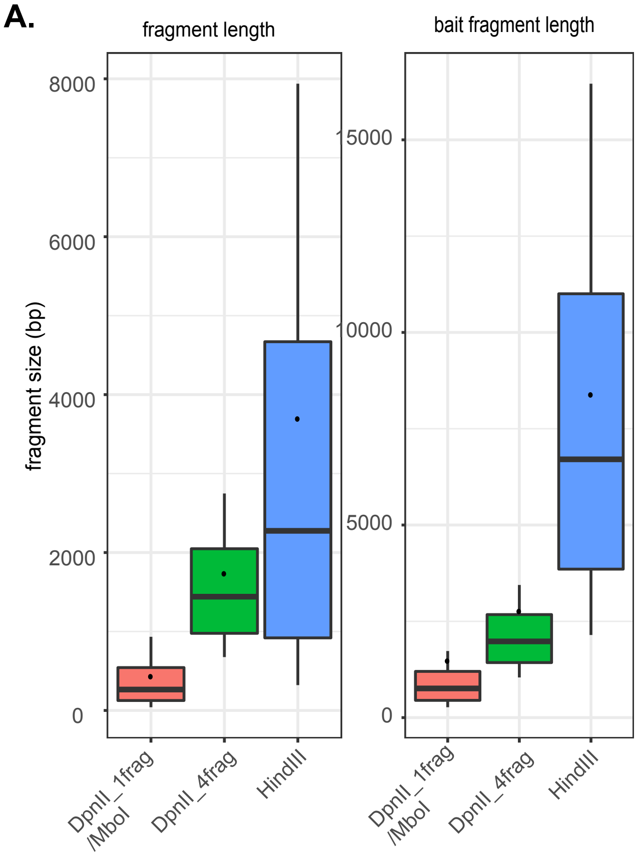


**B.**

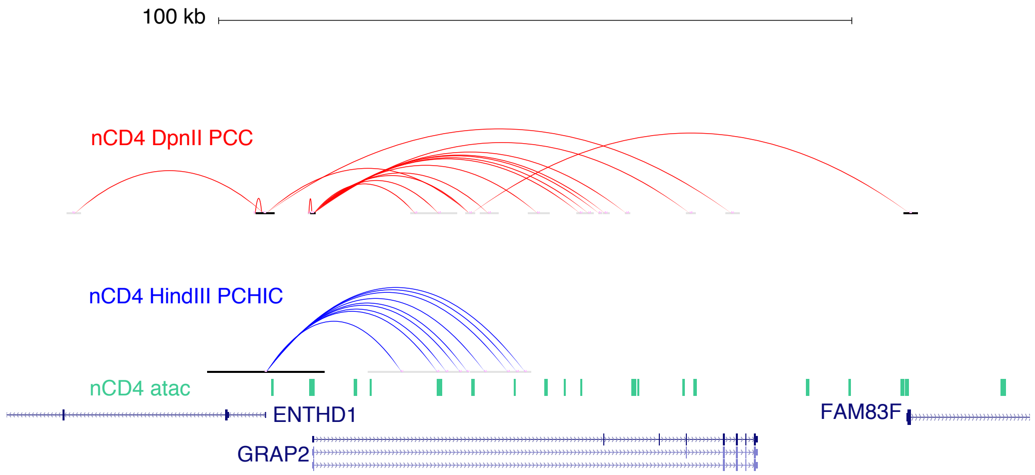

### supfigure3

**Supplemental Figure 3**

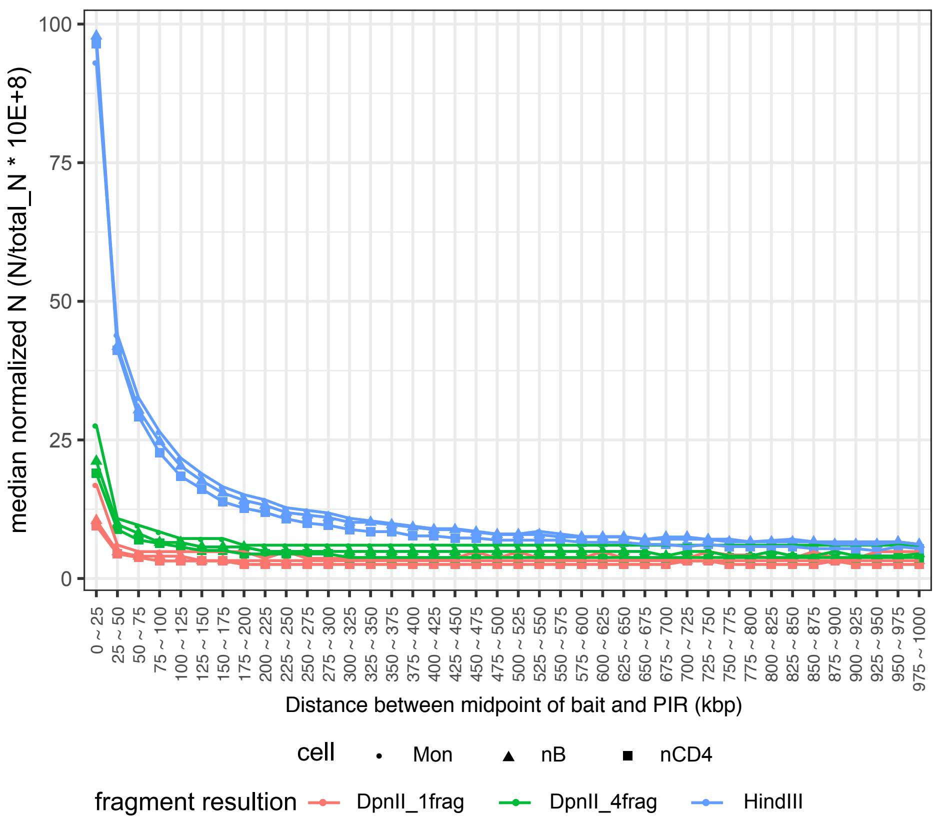

### supfigure4

**Supplemental Figure 4**

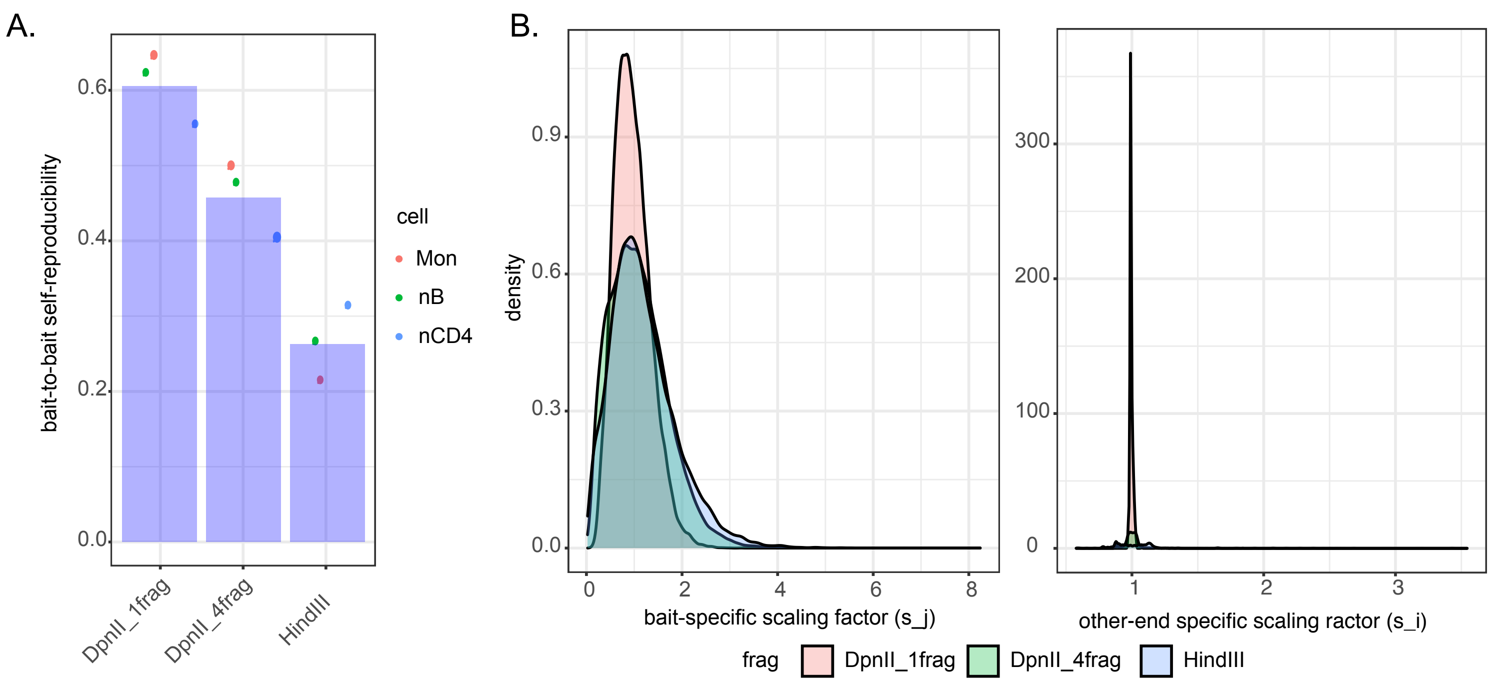

### supfigure5

#### Supplemental Figure 5

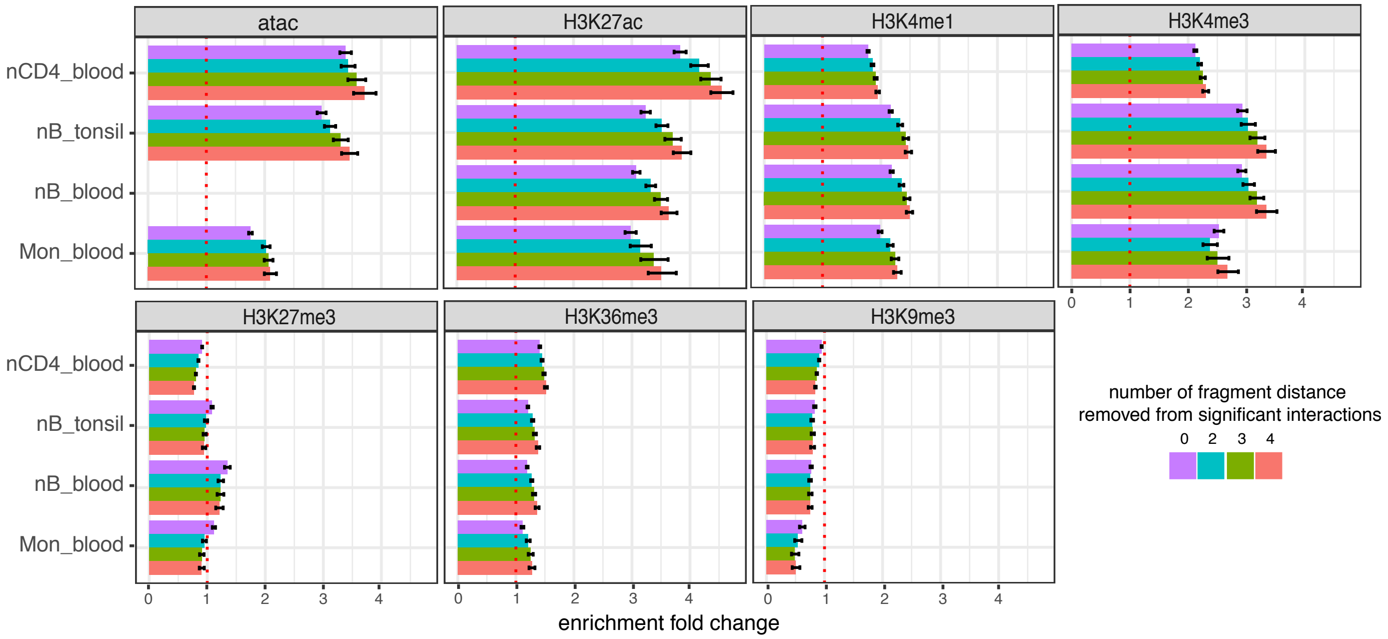

### supfigure6

#### Supplemental Figure 6

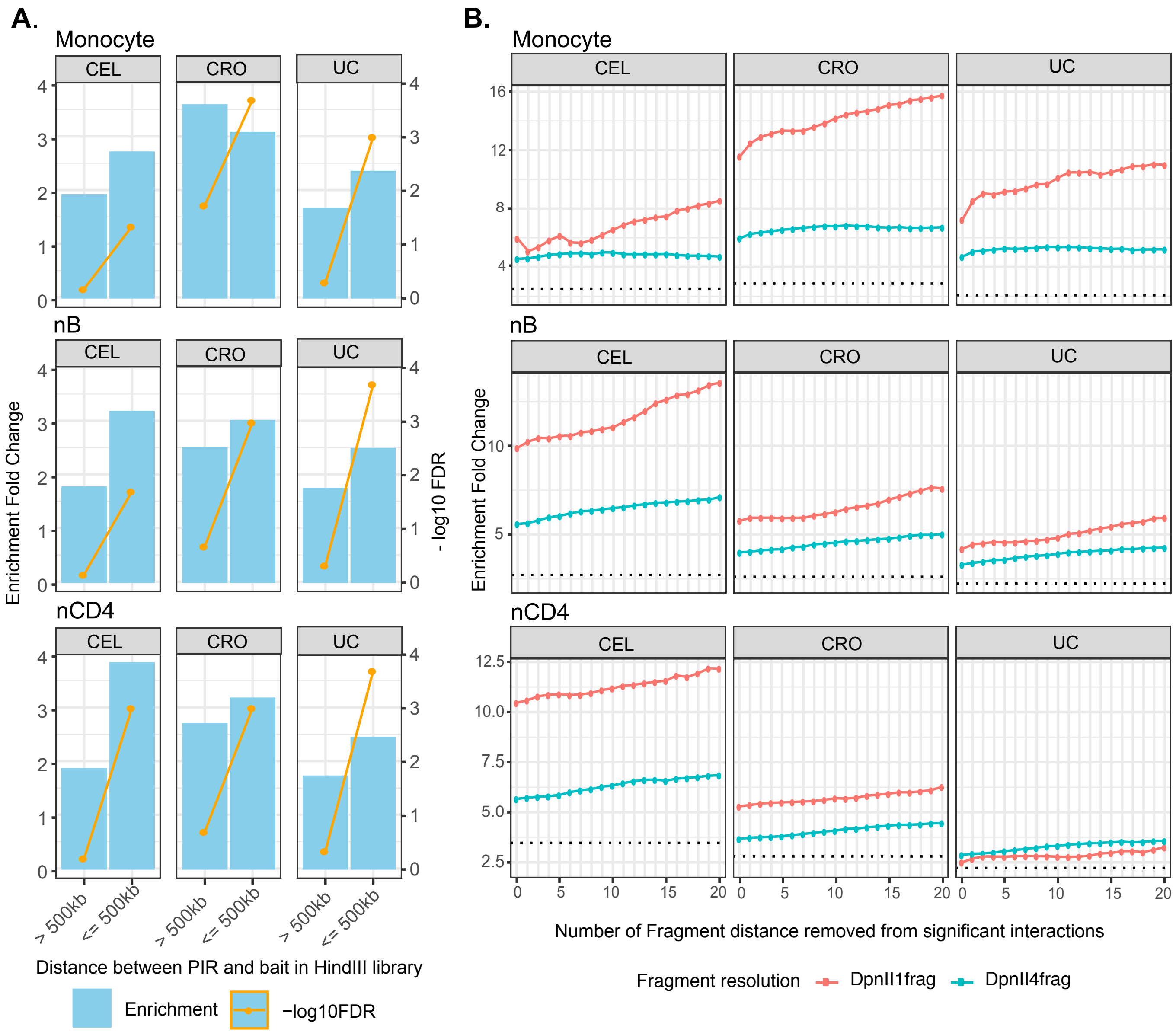
